# Chromosome-level genome assembly of the Endangered scaly-sided merganser *Mergus squamatus* with insight into its demographic history

**DOI:** 10.64898/2026.01.29.702634

**Authors:** Joshua J. Wright, Heleen De Weerd, Alexander C. Lees, Kirsty J. Shaw, Sarah M. Griffiths

## Abstract

The scaly-sided merganser, *Mergus squamatus*, is an Endangered piscivorous duck which has been declining since the late 1900s due to habitat loss, over-hunting, and climate change. Despite being a species of global conservation concern and subject to ex- and in-situ conservation efforts, genomic research has been limited, hindering our understanding of its population genetic status and evolutionary history. In this study, we present the first fully annotated, chromosome-level genome for the scaly-sided merganser, generated using Oxford Nanopore long reads, Illumina short reads, and Hi-C sequencing. The final assembly spans 1.1 Gb across 307 scaffolds, 64 of which are anchored into 35 chromosomes, covering 99.5% of the genome. The assembly shows high contiguity (N50 = 84.3Mb) and completeness, with a Benchmarking Universal Single-Copy Ortholog (BUSCO) score of 98%. Repeat sequences comprise 9.55% of the genome. Homology-based gene annotation identified ∼15,200 protein-coding genes. A complete 16,624 bp mitochondrial genome was also assembled and annotated. Synteny analysis revealed strong chromosomal conservation across the wider Anatidae family, with evidence of lineage-specific rearrangements. Pairwise Sequential Markovian Coalescence modelling indicates recent stability in the effective population size of the species, with past declines coinciding with Pleistocene glacial cycles. Our high-quality genome provides an essential resource for conservation genomic and evolutionary studies of the scaly-sided merganser, supporting ongoing efforts to manage and protect this threatened species.

## Introduction

High-quality whole genome assemblies are transforming our understanding of evolutionary processes in non-model species and providing new insights to inform applied conservation management (Saremi et al. 2019). Genomic studies on threatened species show that small populations are susceptible to genomic erosion and accumulation of deleterious mutations (Abascal et al. 2016; Dussex et al. 2023), leading to severe impacts including reduced fitness, mutational meltdown, and ultimately entrapment in an extinction vortex (Gilpin and Soulé 1986; von Seth et al. 2021), as well as loss of adaptive potential. The fine-scale resolution offered by a whole genome datasets can measure genetic load, identify deleterious variants, quantify adaptive genetic variation, pinpoint genomic regions under selection, link phenotypic traits to genes, and elucidate fine-scale population structure, among many other applications (Allendorf et al. 2010; Lawson et al. 2012; Visscher et al. 2017; Vitti et al. 2013). Furthermore, as each whole genome contains an independent lineage, and with it, many independent loci, this data offers increased statistical power to infer historical demographic processes and assess genetic risk (Schraiber and Akey 2015). This is especially pertinent when samples may be rare, as in many endangered species, allowing the maximum information possible to be extracted from a single sample. This highlights the growing importance of genomic approaches for informing conservation strategies in endangered species

*Mergus squamatus* (scaly-sided mergansers) are piscivorous migratory ducks that breed in fast-flowing stream systems in the Taiga zone of East Asia, migrating south to freshwater and saltwater habitats for the winter (Zeng et al. 2017). It is one of only two riverine landscape specialists in the merganser family (Buckton and Ormerod 2002) and an indicator species of high-quality upland riverine ecosystems (Xu et al. 2021a). Historically, *M. squamatus* was widespread as a breeding bird in the riparian forests of Northeast China and Southeast Russia (BirdLife 2025), but over the past three decades has declined dramatically, owing to threats ranging from climate change to over-hunting and habitat loss (BirdLife 2025; IUCN 2025). The species is now restricted to two key locations: the Silkhote-Alin mountains of Russia and the Changbai mountains of China (Xu et al. 2021a; Xu et al. 2021b), and is classified as Endangered on the International Union for the Conservation of Nature (IUCN) Red List (BirdLife 2025; IUCN 2025). A high-quality reference genome for *M. squamatus* would unlock a number of powerful genetic tools for assessing demographic history, genetic diversity, adaptive potential, and genetic load, helping to inform future conservation management both *in situ* and *ex situ* (Solovyeva and Vartanyan 2019).

Iridian Genomes recently deposited a highly scaffolded *M. squamatus* genome on the National Center for Biotechnology Information (NCBI) database (unpublished 2025; Genbank accession: GCA_051419615.1); however, this study will provide the first fully annotated and functional chromosome-level genome assembly for this species. This genome, compiled using Oxford Nanopore long-read, Illumina paired-end short-read, and Hi-C sequencing data, can be used as a reference genome to enable further genomic work on the species, guiding evidence-based conservation strategies and providing greater understanding of *Mergus* evolution.

## Methods

### Sample collection

*Mergus squamatus* liver and heart tissues were obtained from a freshly deceased adult male individual deposited at the Cambridge Museum of Zoology, UK, in 2023 from a private collector (UMZC: uncatalogued). Samples were immediately placed on dry ice during transportation and then stored at -80°C before genomic DNA extraction.

### DNA extraction and sequencing

DNA was extracted from liver samples using the DNeasy Blood & Tissue Kit (catalog number 69504, Qiagen, Germany). This DNA was subsequently used for Illumina paired-end short-read and Hi-C sequencing. This extraction method proved inadequate for long read sequencing due to fragmentation, so ammonium acetate extractions (Nicholls et al. 2003) were used to obtain suitable materials for this. DNA obtained from both extraction types was tested for purity (260/280 nm and 260/230 nm absorbance ratios) with a Nanodrop 2000 Spectrophotometer (Thermo Fisher, USA) and concentration was measured using a Qubit™ dsDNA Broad Range assay kit (Thermo fisher, catalog: Q33231); DNA was also run on a 1% agarose gel with 1 Kb Hyperladder (catalog number: BIO-33053, Meridian Biosciences, UK) to examine the molecular weight. DNA extracted for long-read sequencing was evaluated on a TapeStation 4200 (Agilent, USA) to ensure average fragment length was ≥30 kb to maximize adequate sequencing. A single Hi-C library preparation was carried out using the Epitect Hi-C kit (catalog 59971, Qiagen, Germany), according to the manufacturer’s instructions and sent to Novogene Europe to be sequenced on the NovaSeq PE150 platform. Two high-molecular weight DNA extracts were sent to Novogene Europe with which five 2×150 base pair (bp) paired-end (PE) libraries were created and sequenced on the NovaSeq 6000. The DNA from the ammonium acetate extraction was also sent to Novogene to be sequenced on the PromethION platform utilizing an insert size of 30 Kb.

### Data quality

The Illumina reads were quality controlled using Trimmomatic V0.38 (Bolger et al. 2014) with the functions: LEADING: 3, TRAILING: 3, SLIDINGWINDOW: 4:20, and MINLEN: 50 to produce a dataset in which the reads surpassed a Phred score of Q20 and were a minimum length of 50 bp.

The Nanopore data were filtered and trimmed using NanoPack2 (De Coster and Rademakers 2023), using NanoPlot V1.40.2 to check the quality of the reads and Chopper V0.8.0 to filter and trim the reads. After visualizing 3’ and 5’ ends of a subset of Nanopore reads, a 27bp head crop and 38bp tail crop was implemented. All reads were trimmed to a mean Q-score of 15 and a minimum length of 500 bp.

### Genome assembly and polishing

Genome assembly was completed using Flye v2.8.1 (Kolmogorov et al. 2019) using the parameters: --nano-hq, --asm-coverage 60, --genome-size 1.2G. Genome size estimation was obtained through the use of Genomes on a Tree (Challis et al. 2023). The genome was then polished firstly using one round of Medaka v1.11.0 (ONT, UK) and subsequently using the Illumina short reads with NextPolish (Hu et al. 2020) following Luan et al. (2024). The Benchmarking Universal Single-Copy Orthologs (BUSCO) (Simão et al. 2015) tool was used at each stage to assess genome completeness.

To further improve the genome assembly and overall completeness, Hi-C data was used to identify genomic regions that are in close physical proximity within the nucleus. This information was then used to scaffold the assembly to chromosome scale. HiCExplorer V3.7.2 (Wolff et al. 2020) was used to quality check and format the Hi-C data. YaHS V1.2a.1 (Zhou et al. 2023) was then used to map the Hi-C reads onto the genome, aiding in misassembly detection and correction to create an updated assembly which was again passed through BUSCO. Decontamination of the assembled genome then took place using Tiara v1.0.3 (Karlicki et al. 2021), a deep-learning tool that removes prokaryotic and organellar DNA from the genome.

### Manual curation

Treeval v1.2.3 (Pointon et al. 2025) was implemented to analyze and create the files required for manual curation. Manual curation was then performed in PretextView v1.0.4 (Harry 2025) to identify and correct misjoins. JBrowse2 (Diesh et al. 2023) was used to visualize the closely related congeneric Brazilian merganser (*Mergus octosetaceous*) and the model organism Zebra Finch (*Taeniopygia guttata*) to guide the rearrangement of the *M. squamatus* genome and to appropriately label the sex chromosomes.

For the final iteration of the genome assembly, completeness was evaluated by mapping short-reads back to the assembly using the Burrows-Wheeler Aligner (BWA; (Li and Durbin 2009) and processing the alignments using Samtools (Danecek et al. 2021) to assess depth and coverage of the genome.

### Mitochondrial genome assembly and annotation

The program MitoFinder v1.4.2 (Allio et al. 2020) was used to identify the mitochondrial genome sequence from the Illumina short-reads, using the NCBI *M. squamatus* mitochondrion sequence HQ833701.1 as reference (Liu et al. 2012). Two iterations of this program were used: the first to identify contigs containing mitochondrial sequences, and the second to refine the annotation using the most suitable contig. The resulting mitogenome was then visually polished in Geneious Prime v2024.0.5 (https://www.geneious.com).

Once obtained, the control region for this mitogenome and HQ833701.1 (Liu et al. 2012) were then extracted and aligned against the five control region haplotypes found across wild populations in Russia and China (Shen et al. 2024b; Solovyeva and Pearce 2011) (accession numbers: HM639863.1, HM639866.1 HM639864.1 HM639865.1, PP828705.1) using Geneious Prime’s Clustal Omega alignment v1.2.3 (Sievers et al. 2011) plugin. Sequence variation among the haplotypes was highlighted within the multiple sequence alignment view within Geneious Prime.

### Repeat annotation

Repeat sequences in the *M. squamatus* genome were annotated *de novo*. RepeatModeler2 v2.0.7 (Flynn et al. 2020), which incorporates RECON, RepeatScout and LTRHarvester, was used to build a database of repeat sequences within the genome. RepeatMasker v4.2.1 (Smit et al. 2015) was then used to extract and annotate these repeat regions.

### Genome annotation – structure and function

Gene annotation was conducted using the homology-based tool LiftOff v1.6.3 (Shumate and Salzberg 2021). Reference protein sequences and whole genomes were downloaded from NCBI for five other Aves species and used as references to identify the most complete annotation through BUSCO analysis: the Mute Swan (*Cygnus olor;* GCF_009769625.2), Mallard (*Anas platyrnchos*; GCF_047663525.1), Tufted Duck (*Aythya fuligulai;* GCA_009819795.1), Red Junglefowl (*Gallus gallus;* GCA_016699485.1) and Zebra Finch (*Taenioygia guttata;* GCA_048771995.1).

### Genome synteny

The conservation of gene order and chromosome structure across the Anatidae was visualized using NGenomeSyn v1.41(He et al. 2023). The above genomes were prepared using SeqKit (Shen et al. 2024a) to retain only autosomes and the Z chromosome. Unlocalized sequences, scaffolds, and the W chromosomes were excluded along with chromosomes below 5 Mbp to ensure parity among each genome sequence. Whole genome alignment was carried out using minimap2 (Li 2018) for species comparison.

### Demographic inference

Pairwise Sequentially Markovian Coalescent (PSMC) v0.6.5 (Li and Durbin 2011) modelling was performed to assess the demographic history of *M. squamatus*. Illumina short-reads were mapped to the SeqKit prepared genome using BWA (Li and Durbin 2009) and a file for the alignment was extracted with a base quality filter using bcftools mpileup (v1.16; (Danecek et al. 2021)) (-Q 30 -q 30). Average coverage of the short reads to the genome was 100x. Sites were then filtered for coverage between 20-120x and a base quality of 30 with vcftools (Danecek et al. 2021) vcf2fq (-d 20 -D 120 -Q 30) and transformed into PSMC files. These coverage filters help prevent low-coverage sites from flattening PSMC demographic curves. PSMC inference was obtained using parameters tested on the sister species *M. octosetaceous* (Granger-Neto et al. 2025) with a coalescent window of 1+1+1+1+1+25*2+4+6, a mutation rate of 5×10^-9^, and a generation time of two years, with 100 bootstrap replicates performed to assess uncertainty.

## Results and Discussion

### Assembly

After trimming and filtering, a total of 100.3 Gb of Oxford Nanopore Technology (ONT) long-reads with a mean read length of 9,236.3 bp and a read quality of 17.4 were obtained, a total of 171 Gb of short reads and 139.3 Gb of Hi-C sequence data. Flye v2.8.1 (Kolmogorov et al. 2019) produced a draft assembly of 1.1 Gb consisting of 607 contigs. As Flye conducted an initial polishing step, further polishing with Medaka & NextPolish did not affect contiguity, however they greatly reduced the number of Ns within the assembly (Initial Flye: 2,400 Ns; Flye+Medaka: 800 Ns, Flye+Medaka+NextPolish: 795 Ns). The utilization of HiCExplorer and YaHS improved the genome contiguity from 607 contigs to 420. Out of the 464,299,476 sequenced HiC reads, there were 59,450,352 Hi-C contacts (12.8%) found and 294,027,555 unique high quality and mappable pairs (63.33%). While these reads improved the orientation of the genome and significantly increased assembly contiguity of the genome table, they also increased the number of Ns in the genome from 795 to 40,395. Tiara further refined the assembly by removing prokaryotic contamination and removing organellar DNA. After manual curation, which focused on correcting misjoins, the final iteration produced the most contiguous assembly, including the largest contig recovered from all versions. The final assembly had an assembly size of 1.1Gb, consisting of 307 contigs, with an N50 of 84,345bp, a GC content of 41.55%, and containing 43,595 Ns (Table 1). The final assembly is a highly contiguous chromosome level assembly of *M. squamatus*, with a total of 35 chromosomes ranging from 0.3-205 Mb in size and covering 99.5% of the whole genome (Figure 1).

**Table 1.** Assembly data and metrics for each step of the draft genome assembly including polishing, Hi-C scaffolding, decontamination, manual curation and final assembly.

| Assembler | Total length | Contig number | Largest contig | N50 length | N50 Number | L90 Length | L90 Number | GC (%) | Ns |
| --- | --- | --- | --- | --- | --- | --- | --- | --- | --- |
| Flye | 1,096,031,551 | 607 | 83,089,125 | 15,125,247 | 16 | 2,831,945 | 75 | 41.56 | 2400 |
| Flye+Medaka | 1,096,122,031 | 607 | 83,098,387 | 15,126,319 | 16 | 2,832,064 | 75 | 41.55 | 800 |
| Flye+Medaka+NextPolish | 1,095,963,894 | 607 | 83,086,067 | 15,123,680 | 16 | 2,831,844 | 75 | 41.55 | 795 |
| Assembly+HiC | 1096,003,494 | 420 | 204,650,930 | 84,345,675 | 4 | 15,192,112 | 18 | 41.55 | 40,395 |
| HiC Assembly+Tiara | 1,095,427,760 | 323 | 204,650,930 | 84,345,675 | 4 | 15,983,245 | 17 | 41.55 | 40,195 |
| Post-curation | 1,095,431,160 | 307 | 205,034,483 | 84,345,675 | 4 | 15,192,112 | 18 | 41.55 | 43,595 |

Post-curation is the final assembly used for downstream genomic analysis. Manual curation was carried out using PreText View (Harry, 2025) to identify and correct misjoins within the assembly

**Figure 1.**
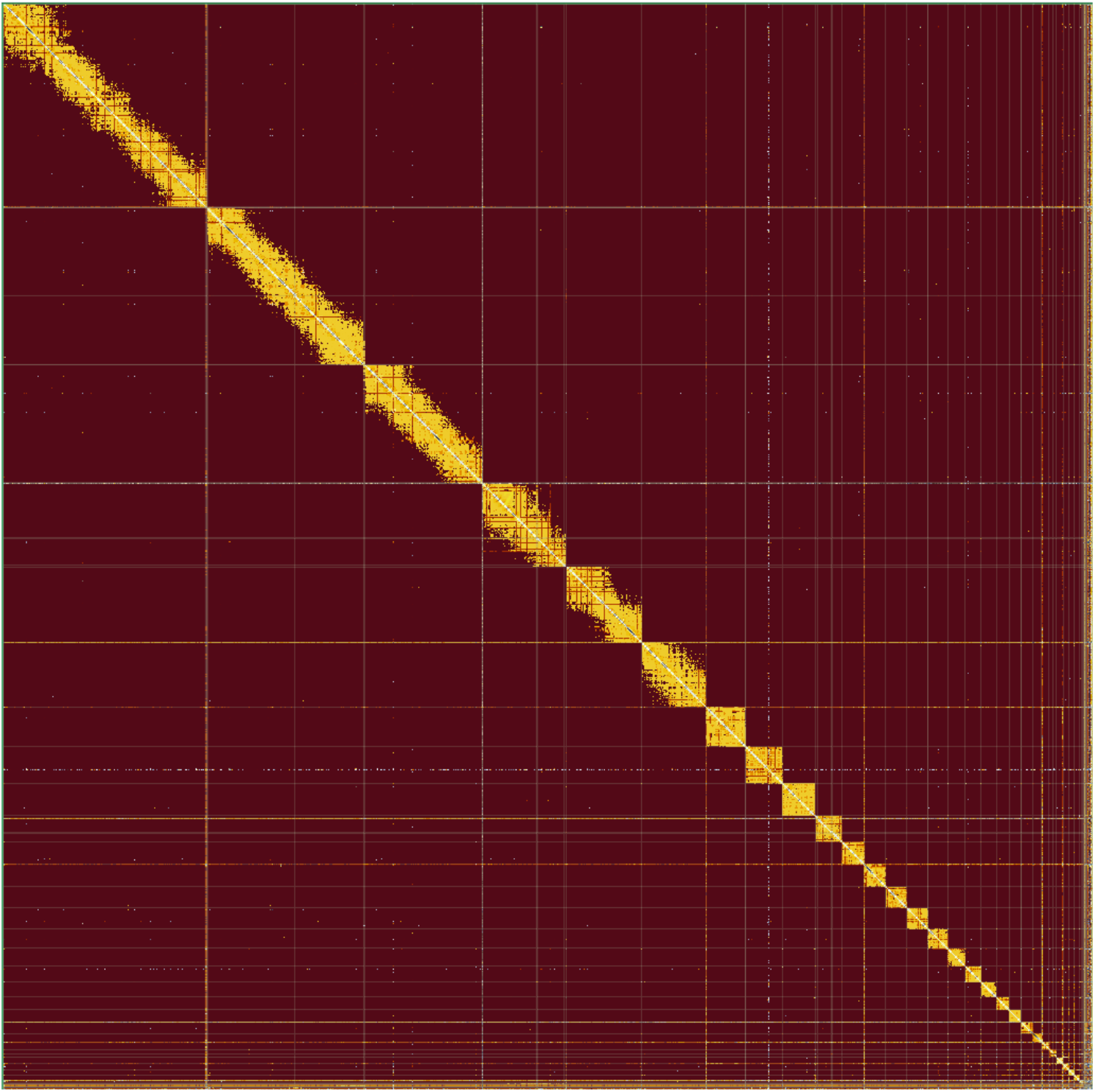
Chromosome heatmap of *Mergus squamatus*. Chromosomes are represented in size order from left to right, top to bottom

This is the first chromosome-level genome assembly available for *M. squamatus*, achieving a much more contiguous genome (Table 1) than the existing *M. squamatus* assembly, which comprised of 463,241 scaffolds (unpublished 2025; Genbank accession: GCA_051419615.1). The size of genome produced in this study is consistent with other members of the Anatidae, such as the mallard (*A. platyrhynchos*) at 1.3 Gb (GCA_047663525.1), and the Brazilian merganser (*M. octosetaceous*) at 1.2 Gb (GCA_036873955.1). The GC content of this assembly (41.55%) is also in line with other Aves, for which GC content typically falls between 40-45% (Zhang et al. 2014). In this genome, 34 autosomes and the Z sex chromosome were identified. This is close to the expected karyotype for Anatidae (n=40; (Beklemisheva et al. 2024). As is characteristic of birds, several microchromosomes were present; these typically require higher sequencing resolution and are among the most challenging to assemble and curate (Peona et al. 2018).

### Genome completeness

BUSCO scores suggest high assembly completeness. Using the aves_odb12 database (Tegenfeldt et al. 2025) of 6,251 single-copy orthologs, 98% were present in the final chromosome assembly (Table 2). This is slightly lower than the completeness of the sister species *M. octosetaceous* (98.9%; (Granger-Neto et al. 2025) but is still highly complete.

**Table 2.** BUSCO analysis results of *Mergus squamatus* genome.

| Category | Number | BUSCO (%) |
| --- | --- | --- |
| Complete BUSCOs | 6128 | 98.00 |
| Complete and single-copy BUSCOs | 6099 | 97.57 |
| Complete Duplicated BUSCOs | 29 | 0.46 |
| Fragmented BUSCOs | 44 | 0.70 |
| Missing BUSCOs | 79 | 1.26 |
| Total BUSCO groups searched | 6251 |  |

Using the BWA-MEM algorithm (Li and Durbin 2009), the alignment rate of all short-reads against the genome was 99.02%, and the breadth of coverage was 99.97%. The mean sequencing depth across the assembly was approximately 153x, indicating remarkably high coverage and concordance between the short-reads and the assembled genome, with almost complete alignment and coverage.

### Repeat annotation

Using RepeatModeler2 and RepeatMasker, approximately 104.6 Mb of repeat sequences were identified in the genome, accounting for 9.55% of the whole genome. Among the repeat sequences identified, short interspersed nuclear elements (SINEs) accounted for 0.04% of the genome, long interspersed nuclear elements 5.44%, and long terminal repeats (LTRs) 0.97% (Table 3). For tandem repeats, satellite DNA composed 0.01% and simple repeats amounted to 2.21%.

**Table 3.** Annotation of repeated sequences.

| Repeat type | Length (bp) | Percentage of Assembled Genome (%) |
| --- | --- | --- |
| DNA transposons | 1,412,690 | 0.13 |
| Tandem repeats | 24,180,385 | 2.22 |
| Small RNA | 119,897 | 0.01 |
| LINE | 59,587,980 | 5.44 |
| SINE | 468,901 | 0.04 |
| LTR | 10,589,094 | 0.97 |
| Unknown | 8,148,347 | 0.74 |
| **Total** | **104,507,294** | **9.55** |

LINE: long interspersed nuclear elements; SINE: short interspersed nuclear elements; LTR: long terminal repeats

The proportion of repeat sequences in the *M. squamatus* genome (9.55%) is relatively low for the Anatidae. For comparison, *Cygnus olor* contains 7.71% repeats (Chong et al. 2023), *M. octosetaceous* contains 18.9% (Granger-Neto et al. 2025), *Aythya fuligula* contains 13% (Mueller et al. 2021), and *Anas platyrhynchos* contains 15% (Mueller et al. 2021).

### Gene annotation – structure and function

Using homology-based annotation, several gene sets were produced, and the most informative set was selected for analysis. On average, 14,732 genes were identified across all reference species, with ∼11 exons per gene. The average coding DNA sequence (CDS) length was 1,773 bp, while mean exon and intron lengths were 274 bp and 3,021 bp, respectively (Table 4).

**Table 4.** Prediction of protein-coding genes.

| Methods/Tools |  | Gene number | Average exons per gene | Average Length (bp) |  |  |  |
| --- | --- | --- | --- | --- | --- | --- | --- |
|  |  |  |  | Transcript | CDS | Exon | Intron |
| Homology | Liftoff: *Anas platyrhynchos* | 15,201 | 11.4 | 36,868 | 1,857 | 320 | 3,196 |
|  | Liftoff: *Aythya fuligula* | 14,519 | 11.4 | 35,273 | 1,834 | 244 | 3,118 |
|  | Liftoff: *Cygnus olor* | 14,990 | 11.4 | 36,227 | 1,840 | 306 | 3,135 |
|  | Liftoff: *Gallus gallus* | 14,752 | 10.5 | 33,846 | 1,761 | 338 | 3,199 |
|  | Liftoff: *Taeniopygia guttata* | 13,427 | 10.1 | 33,707 | 1,733 | 319 | 3,330 |

BUSCO analysis on each data set showed a difference between number of genes annotated and the completeness of the gene sets (Table 5), with clear evidence of distance decay between *M. squamatus, G. gallus*, and *T. guttata*. LiftOff between the *M. squamatus* and *A. platyrhynchos* was the most complete and functionally annotated, so this annotation was used for downstream analysis.

**Table 5.** BUSCO analysis on predicted genesets.

| Geneset | Complete | Single | Duplicated | Fragmented | Missing |
| --- | --- | --- | --- | --- | --- |
| Homology/Liftoff: *Anas platyrhynchos* | 95.90% | 43.50% | 52.40% | 0.30% | 3.80% |
| Homology/Liftoff: *Aythya fuligula* | 94.40% | 63.30% | 31.10% | 0.80% | 4.80% |
| Homology/Liftoff: *Cygnus olor* | 89.30% | 41.60% | 47.70% | 1.90% | 8.80% |
| Homology/Liftoff: *Gallus gallus* | 73.40% | 26.00% | 47.40% | 4.30% | 22.30% |
| Homology/Liftoff: *Taeniopygia guttata* | 59.50% | 26.80% | 32.70% | 6.90% | 33.60% |

LiftOff is a homology-based annotation tool, so the quality of the gene set produced is based strongly off the original genome’s annotation. Therefore, homologous annotation should be performed using high quality references to avoid propagating existing errors (Yandell and Ence 2012). The gene set produced using LiftOff with the *A. platyrhynchos* annotation produced the highest quality annotation (BUSCO 95.90%), with many high-quality genomes having annotation BUSCOs between 94-98% (Simão et al. 2015). While the annotation BUSCO for this assembly is marginally lower than the assembly for this genome, this is a common occurrence during annotation (Manni et al. 2021).

### Mitochondrial genome assembly and annotation

We assembled a complete mitochondrial genome of *M. squamatus*. This mitogenome is 16,624 bp in length with 49.6% GC content (Figure 2). All 37 expected vertebrate mitochondrial genes were annotated. Only minor differences were found between this mitogenome and the reference mitogenome used to map and assemble our reads against (HQ833701; (Liu et al. 2012), which was 29 bp smaller at 16,595 bp and with a GC content of 48.7%.

**Figure 2.**
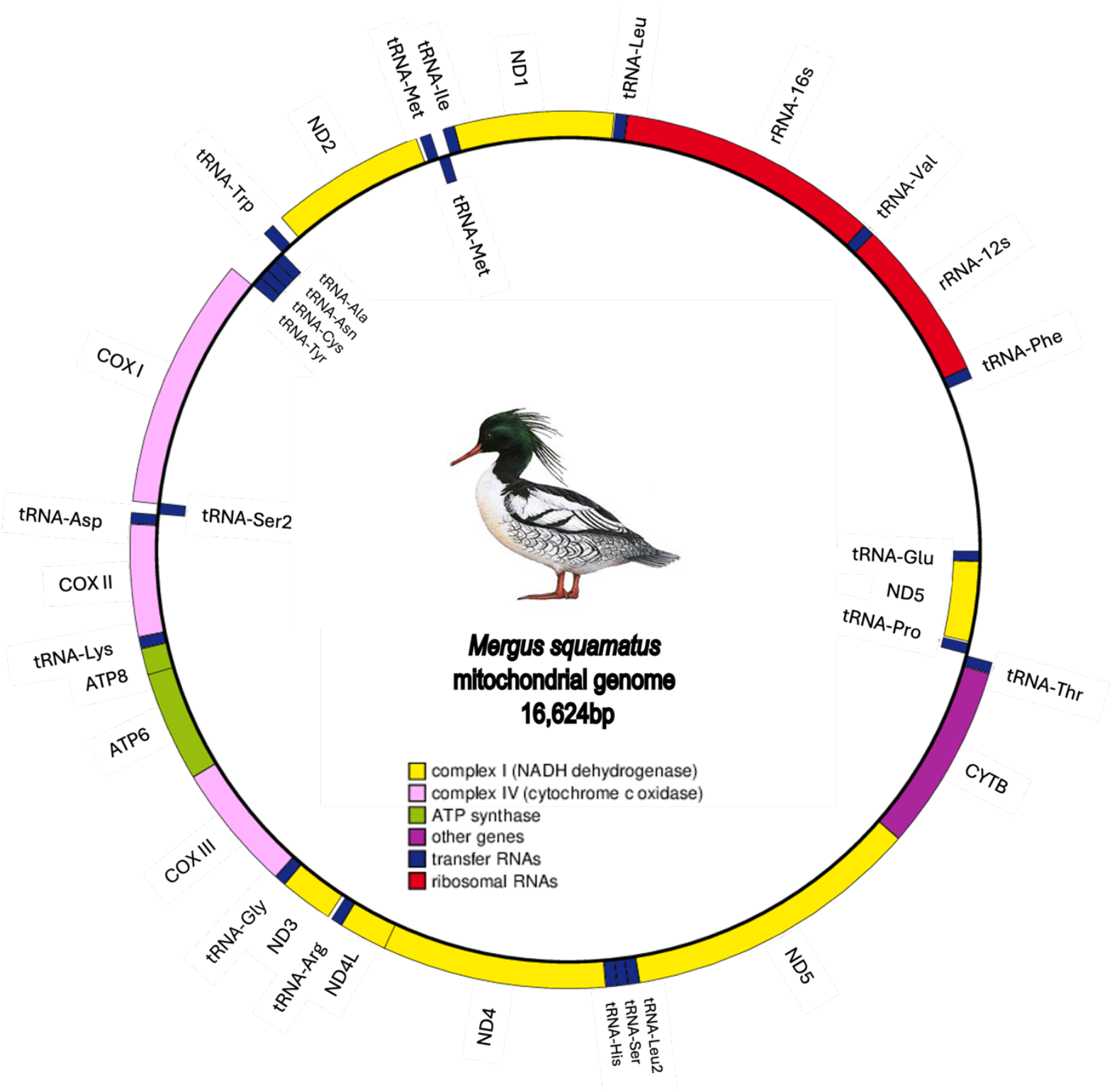
Complete annotated mitochondrial genome of *Mergus squamatus*

Unlike the closely related *M. octosetaceous*, the ND3 gene was found as a whole unit rather than two subunits (Granger-Neto et al. 2025), the separation of which is seen in other Anatidae species (e.g. *Aythya baeri;* (Liu et al. 2021). Studies of the *M. squamatus* mitochondrial control region have identified only six haplotypes in the wild (including the reference mitogenome HQ833701) (Liu et al. 2012; Shen et al. 2024b; Solovyeva and Pearce 2011). The analysis carried out here shows our mitogenome control region haplotype is identical to one of the previously discovered haplotypes.

### Syntenic analysis

Our syntenic analysis showed widespread genomic conservation across the Anatidae (Figure 3). Strong conservation of synteny can be seen across all autosomal chromosomes and the Z chromosome, with syntenic blocks covering the whole chromosomes. There is also however, some evidence of inversion events happening within macrochromosomes across the genomes. All Anatidae species evaluated share the same karyotype (n=40). However, *M. octosetaceous* has a larger chromosome 9, resulting from the fusion of this chromosome and a microchromosome.

**Figure 3.**
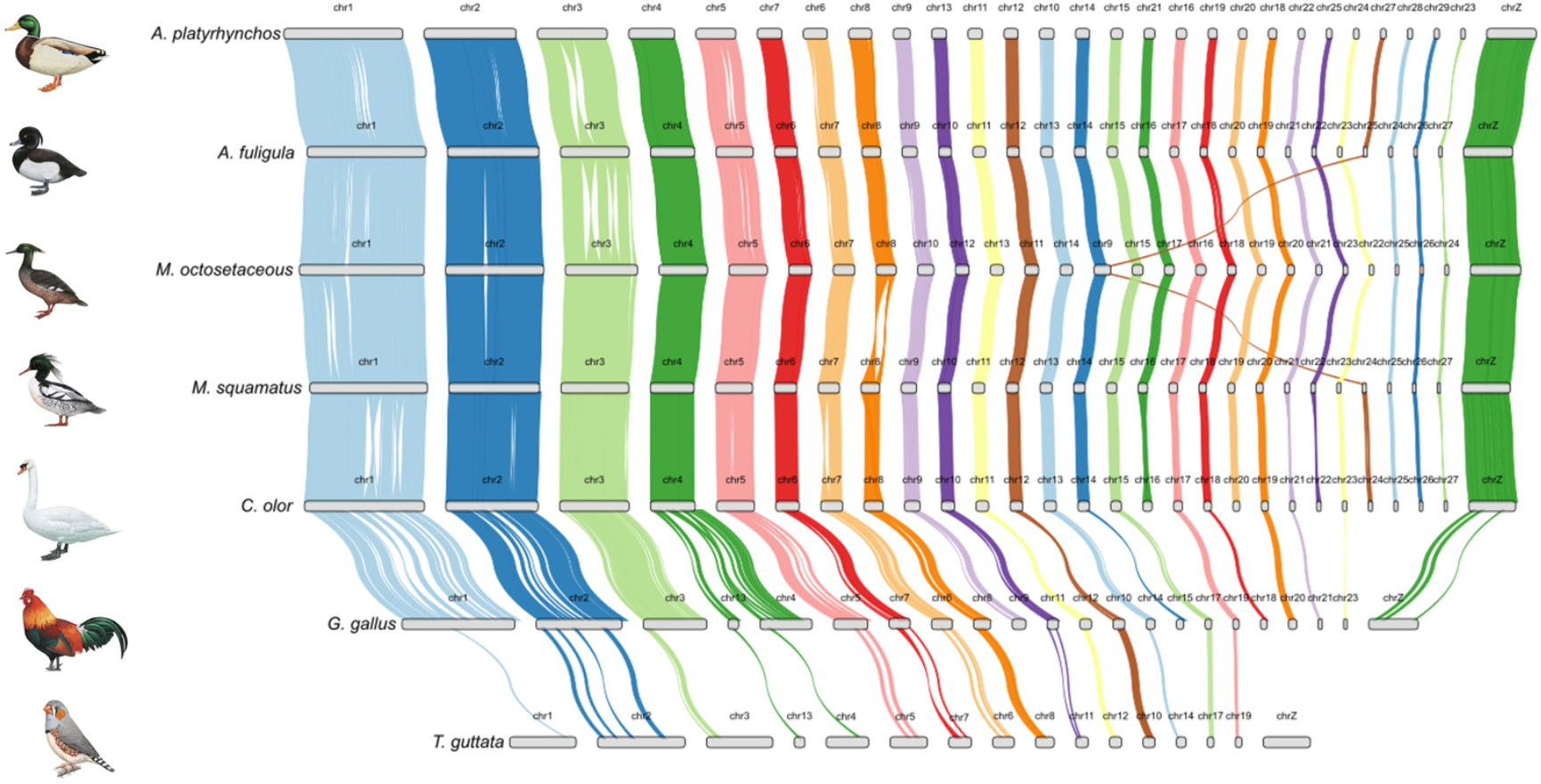
Syntenic analysis of chromosomes from representative Anatidae species with *Gallus gallus* and *Taeniopygia guttata* as outgroups as used in gene annotation. Colored blocks represent homologous chromosomes, with colors conserved across species to indicate shared ancestry. Inversions are visualized as”hourglass” shapes between syntenic blocks to show where homologous chromosomes are reversed between species. Chromosome fusions are indicated by the merging of colored blocks across species i.e chr25 and chr24 of *Aythya fuligula* and *Mergus squamatus* respectively and chr9 of *Mergus octosetaceous*.

This fusion of chromosomes can be seen in other species within the Anatidae, such as *Oxyura jamaicensis* which has a fused chromosome while still retaining the karyotype n=40 (Beçak et al., 1973), suggesting the conservation of this karyotype among this family happens through structural changes within the genomes (Beklemisheva et al. 2024)

### Demographic history

Demographic history analysis of the *M. squamatus* with PSMC revealed that the effective population size (N_e_) has been declining gradually over the past 1.2 millrion years (Figure 4). This decline accelerated during the glacial cycles of the Mid-Pleistocene before plateauing during the Eeminan interglacial period. Population size recovered slightly ∼40kya before a pronounced decline as temperatures dropped during the Last Glacial Maximum (∼19-27 kya) and then remained at a plateau until the inference limit of 10 kya.

**Figure 4.**
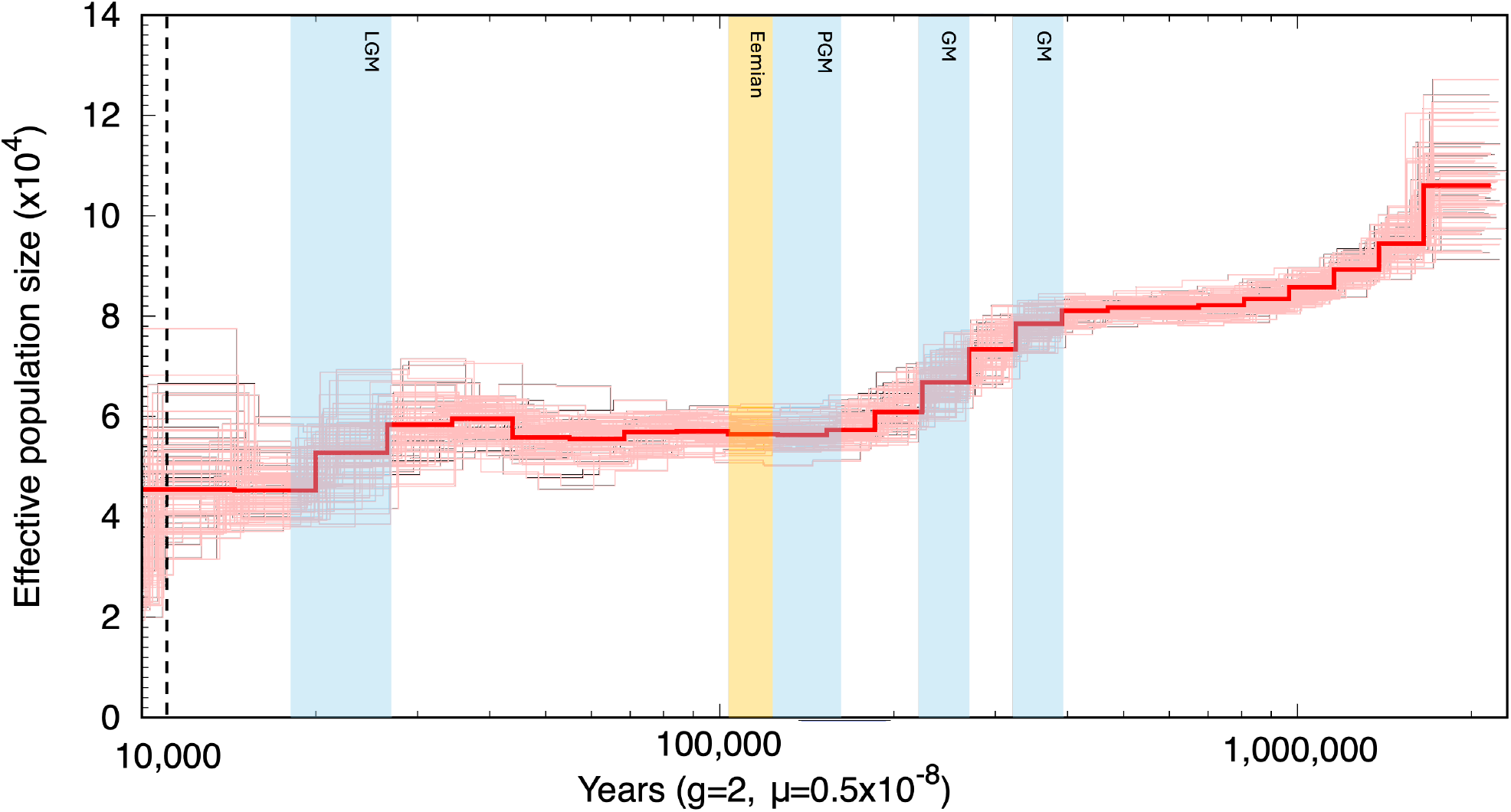
Demographic reconstruction of *Mergus squamatus* based on autosomal chromosomes with PSMC. Estimates used a generation time of 2 years and a mutation rate of 0.5×10^-8^ substitutions per site/generation as optimised by Granger-Neto et al., 2025. Bootstrap replicated (N=100) are shown by the lighter red lines. Glacial maxima are presented with light blue bars whereas the Eemian warm period is represented by the light orange with names listed on the bars. LGM: Last glacial maxima, PGM: Penultimate glacial maxima, GM: Glacial maxima. Glacials and interglacials were modified from Webster et al. (2009). The hashed line represents the transition between the Pleistocene and Holocene epoch. Yearsarerepresentedon a logarithmic scale ranging from 10,000 to 1,200,000 years ago

In comparison to the Critically Endangered *M. octosetaceous* (Granger-Neto et al. 2025) *M. squamatus* has maintained an overall higher effective population size (N_e_). Around 10,000 years ago, *M octosetaceous* had an N_e_ of approximately 15,000, whereas *M. squamatus* exhibited a substantially larger N_e_ of ∼48,000. This may reflect the historical differences in species ecology, with *M. squamatus* being more widely distributed than the more range-restricted *M. octosetaceous*. However, while this is a helpful estimate of N_e_, rapid population declines that ensued during glacial maxima may also be misinterpreted as populations subdividing or large migration events leading to potential bottlenecks (Bansal and Nichols 2025). Yet, as PSMC establishes this N_e_ model from a single diploid genome it is strongly limited to individual specific variation, where more coalescent events can be found in increased sample sizes (Terhorst et al. 2017).

### Summary

Here we present a highly contiguous (307 scaffolds, 35 chromosome level, 1.1 Gb) and complete (BUSCO score of 95.5%) genome for *M. squamatus*, which has immediate uses in aiding conservation of genetic resources in this Endangered species. As sequencing capacity increases and reference genomes are developed, our ability to explore new conservation genomic avenues become available. This includes the development of novel markers to aid basic conservation management and many more refined analyses, such as gene characterization, which may help us better understand selection pressures on endangered species. Genome Wide Association Studies (GWAS) have become invaluable to the identification and exploratory analysis of gene families essential in key biological processes of threatened species, such as reproduction, behavior and immunity (Brandies et al. 2019). Additionally, this genome may provide uses in comparative genomic research on the evolution of Anatidae and wider Aves in general.

## Data availability

The Mergus squamatus genome sequence is available upon request and will be available through NCBI Bioproject: PRJNA1367167. This will be made available in June 2026. Sequences used to assemble the genome are available through the SRA under the same accession. All code used for to generate data, is available at https://github.com/JWrighty97/SSM_GenomeAssembly-Anno. All other genomes used are referenced and are publicly available on NCBI.

## Acknowledgements

We thank Dr Daniel Field (Cambridge Museum of Zoology) for donating the sample for this work. We would also like to thank Birds of The World for providing the scientific illustrations for each species.

## Author contributions

JJW: molecular laboratory work, bioinformatics, manuscript writing and editing; HDW: bioinformatics support and manuscript editing; ACL & KJS; manuscript editing; SMG: funding acquisition, project conception and manuscript editing. All authors read and approved the manuscript.

